# A *Drosophila* CRISPR/Cas9 toolkit for conditionally manipulating gene expression in the prothoracic gland as a test case for polytene tissues

**DOI:** 10.1101/354258

**Authors:** Nhan Huynh, Kirst King-Jones

## Abstract

Targeting gene function with spatial or temporal specificity is a key goal in molecular genetics. CRISPR-Cas9 has greatly facilitated this strategy, but some standard approaches are problematic. For instance, simple tissue-specific or global overexpression of Cas9 can cause significant lethality or developmental delays even in the absence of gRNAs. In particular, we found that Gal4-mediated expression of UAS-Cas9 in the *Drosophila* prothoracic gland (PG) was not a suitable strategy to disrupt gene expression, since Cas9 alone caused widespread lethality. The PG gland is widely used for studying endocrine gland function during animal development, but tools validating PG-specific RNAi phenotypes are lacking. Here, we present a collection of modular gateway-compatible CRISPR-Cas9 tools that allow precise modulation of target gene activity with temporal and spatial specificity. We also demonstrate that Cas9 fused to the progesterone ligand-binding domain can be used to activate gene expression via RU486. Using these approaches, we were able to avoid the lethality associated with simple GAL4-mediated overexpression of Cas9 in the PG. Given that the PG is a polytene tissue, we conclude that these tools work effectively in endoreplicating cells where Cas9 has to target multiple copies of the same locus. Our toolkit can be easily adapted for other tissues and can be used both for gain-and loss-of-function studies.

## Introduction

It is crucial to have the ability to conditionally manipulate the activity of genes, be it to overcome embryonic lethality of null mutants to study later roles of a given gene, distinguish between cell-autonomous and non-autonomous mechanisms, or to study tissue-specific gene functions. In Drosophila, the standard techniques for conditionally altering gene function have been RNA interference (RNAi) to block or impair gene activity and the overexpression of cDNAs for gain-of-function studies. Most commonly, both RNAi and cDNA expression are temporally controlled via the Gal4-UAS system, resulting in a highly versatile set of tools. However, each of these commonly used components has its limitations and downsides. In particular, RNAi suffers from the frequent occurrence of off-targets, requiring rigorous validation, and often the expression of a target mRNA is only partially blocked. In addition, combining two or more RNAi transgenes to test for synthetic lethality or interaction of pathway components is cumbersome and exponentially increases the risk of non-specific effects. On the other hand, to achieve overexpression of a gene of interest, traditional cDNA overexpression requires the cloning of a full-length cDNA, which may be difficult and time-consuming. Further, in the case of alternatively spliced genes, one usually has to choose which isoform to use for the transgenic cDNA line, which may limit the conclusions that can be drawn from the experiment. It should also be noted that the use of Gal4 itself has its drawbacks. In particular, Gal4 may result in signal amplification due to its strong activation domain, and one has only limited control over how strongly a given cDNA is expressed. Further, UAS-regulated transgenes all show some degree of leakiness, depending on the tissue and developmental time, potentially confounding experimental outcomes (Akmammedov et al., 2017). Similarly, the presence of a second unrelated UAS-transgene may alter phenotypes seen with a single UAS-transgene alone, as both compete for Gal4-binding, which may quench the expression of either transgene. Finally, Gal4 binds non-specifically to endogenous loci, resulting in the up-and down-regulation of hundreds of genes, which may complicate the interpretation of genome-wide gene expression studies (Liu and Lehmann, 2008).

The recent discovery of Clustered Regularly Interspaced Short Palindromic Repeats (CRISPR) and the generation of guide RNA- (gRNA-) dependent Cas9 endonucleases has been quickly adapted by Drosophila researchers (Gratz et al., 2014; Port et al., 2014; Lin et al., 2015) and we now possess a universal and powerful toolkit that can be used for both loss- and gain-of-function studies by using distinct versions of Cas9 (Gupta and Musunuru, 2014; Bier et al., 2018). As such, CRISPR-based techniques are ideal to replace, validate and complement traditional approaches relying on conditionally expressing RNAi or cDNAs. Recent advances in CRISPR-based approaches include codon-optimizations of Cas9, utilizing Cas9 variants as a RNA-guided transcription factors that block or increase target gene transcription, and large-scale transgenic Drosophila gRNA collections launched at Harvard Medical School (https://fgr.hms.harvard.edu/vivo-crispr-0), the German Cancer Research Center in Heidelberg (https://www.crisprflydesign.org/library/) and the National Institute of Genetics in Mishima, Japan (https://shigen.nig.ac.jp/fly/nigfly/).

Our lab investigates signaling pathways that control ecdysone and heme biosynthesis in the larval prothoracic gland (PG), which is part of the larval ring gland (Figure 1). The PG is an endoreplicating tissue that reaches a C-value of 64 by the end of the 3^rd^ instar (L3) (Ohhara et al., 2017) and represents a popular model for studying endocrine function, as multiple checkpoints converge on this gland that dictate whether an upcoming pulse of ecdysone can be produced (Ou et al., 2016). In a recent study, we carried a genome-wide PG-specific RNAi screen, resulting in the identification of ~1,906 genes that where critical for larval development (Danielsen et al., 2016). However, a frequent issue in the follow-up studies was that we could not validate the RNAi-induced phenotypes by independent non-overlapping RNAi lines, either because no such lines existed or because independent lines did not replicate the phenotype. This prompted us to look into CRISPR-based methods that could be used to confirm the RNAi results. However, no studies have addressed whether somatic CRISPR is feasible in the PG, nor have there been any reports on the usage of tissue-specific CRISPR/CAS9 for other commonly studied polytene tissues such as the larval fat body and the salivary glands. Previous studies have established that somatic CRISPR/CAS9 is highly efficient in disrupting genes in a biallelic fashion, however endoreplicating tissues such as the salivary gland contain up to 1024 copies of a gene (Hochstrasser, 1987; Andrew et al., 2000) raising the question as to whether CRISPR/CAS9 would be effective in polytene tissues. Furthermore, our initial attempts to express Cas9 via the most commonly used PG-specific Gal4 drivers resulted in substantial larval lethality, which rendered this approach impractical. We therefore developed several strategies that induced tissue-specific CRISPR/CAS9 without using Gal4.

**Figure 1.**
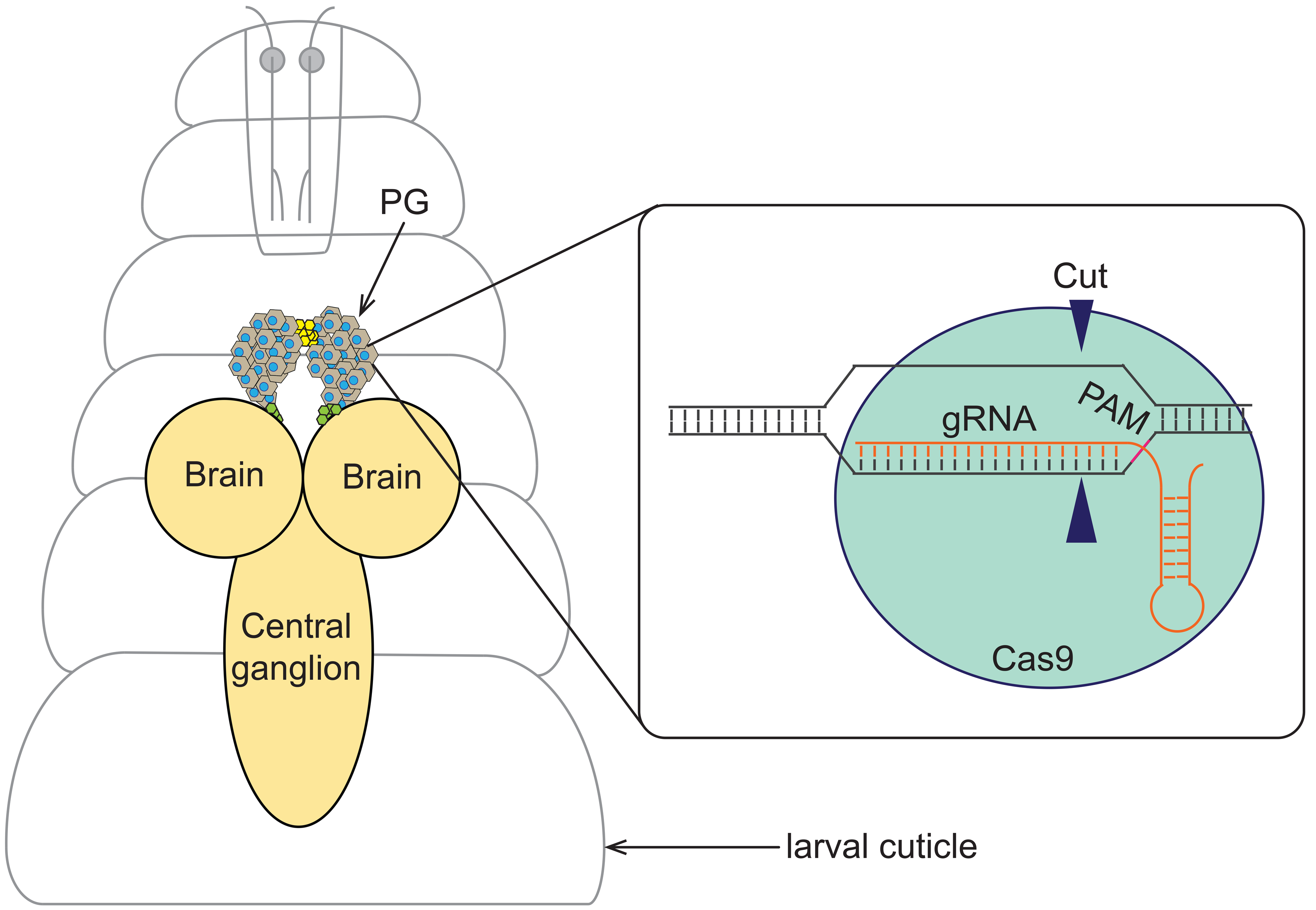
Somatic CRISPR in the *Drosophila* prothoracic gland. In Drosophila larvae, the prothoracic gland (PG) is the principal source for ecdysteroid production. The PG is a part of the ring gland, which also harbours the corpora allata (yellow) and corpora cardiaca (green). PG-specific genome editing via <u>C</u>lustered <u>R</u>egularly <u>I</u>nterspaced <u>S</u>hort <u>P</u>alidromic <u>R</u>epeats (CRISPR) requires the recruitment of CRISPR-associated protein 9 (Cas9: blue) to the target site recognized by the guide RNA (gRNA: orange). Target site cleavage by Cas9 is ensured by the presence of the protospacer adjacent motif (PAM: purple) sequence immediately following the target site. This sequence will direct the cut site of Cas9 to a region of about three nucleotides upstream of the PAM.

Here we present a collection of CRISPR tools designed for tissue-specific genome modification in *Drosophila*, which we refer to as the general Gateway Cas9 (gG-Cas9), the PG-Cas9 and the PG-gRNA vector collections. These new tools have in common that they are not based on Gal4, but rather use enhancer regions to achieve tissue-specific expression of Cas9 or gRNAs. This greatly simplifies the genetics of tissue-specific CRISPR/Cas9, since one only requires a single cross to build the CRISPR/CAS9-gRNA combination, while the Gal4-UAS-based approach requires combining at least three transgenes. Importantly, we show that the lethality associated with Gal4-driven Cas9 can be prevented by several strategies, including tissue-specific expression of gRNAs coupled with ubiquitously expressed Cas9. To accomplish tissue-specific gRNA production, we took advantage of inserting ribozyme sequences, which demonstrates for the first time in Drosophila the that ribozymes can be used to effectively release gRNAs from mRNAs. We also present the first fusion of Cas9 with the Ligand-binding domain of the human progesterone receptor and demonstrate that it is a highly effective tool for achieving both temporal and spatial control over Cas9-mediated gene activation. We evaluated the efficiency of each tool by targeting two well-studied genes acting in ecdysone biosynthesis, *phantom* and *disembodied*. Finally, we provide a general, gateway-based vector collection (gG-Cas9) that allows the quick generation of seven different Cas9-based vectors. These tissue-specific vectors enable the user to i) disrupt target genes of interest, ii) block the assembly of the transcription apparatus near the transcription start site or iii) upregulate the activity of a given gene.

## Results and Discussion

### PG-specific expression of Cas9 via the Gal4/UAS system is toxic

#### a. Classic Gal4/UAS-Cas 9

Current CRISPR/Cas9 tools that conditionally modify gene expression include UAS-Cas9 for GAL4-directed sequence cuts, as well as UAS-dCas9-VPR for gene overexpression, which utilizes a non-cutting version of Cas9 (dead Cas9 = dCas9) fused to the VPR co-activator domain (Port et al., 2014; Lin et al., 2015; Port and Bullock, 2016). Our initial attempts to block gene function in the prothoracic gland (PG) were based on expressing UAS-Cas9.C (the original Cas9 transgene) with *phm22-Gal4* (aka *phm22>Cas9.C* animals), a widely used PG-specific Gal4 driver (Rewitz et al., 2009). However, this approach caused significant lethality, with only ~15% of animals reaching adulthood compared to ~85% in controls (Figure 2A). We then tried another PG-specific Gal4 driver, *spok-Gal4* (= *spok>*), which has overall lower expression levels compared to *phm22>* (Mike O’Connor, personnel communication). This combination resulted in only slightly improved survival rates, with 25% of the population reaching adulthood (Figure 2A). This observation is consistent with previous studies where high expression levels of Cas9 via Gal4/UAS caused toxicity that was independent of the endonuclease activity (Port et al., 2014). The lethality was also observed when we tried different Cas9 versions, namely Cas9.P (codon-optimized for Drosophila) and Cas9.P2 (codon-optimized for human cells) (Port et al., 2014; Xue et al., 2014; Port and Bullock, 2016). UAS-Cas9.P2 was considered to be safer for using the Gal4/UAS approach (Port and Bullock, 2016). Unfortunately, in our hand, *phm22>Cas9.P2* animals, showed only moderately improved survival rates compared to *phm22>Cas9.P* and *phm22>Cas9.C* populations, with only ~50% reaching the third instar stage, and 35% surviving to adulthood (Figures 2 and S1A). Using *spok-Gal4* instead of*phm22-Gal4* as a PG-specific driver did not make a significant difference (Figure S1B). Interestingly, ubiquitous expression of *Cas9.P2 (act>Cas9.P2)* caused no obvious lethality and the majority of the population reaches adulthood, while *act>Cas9.C* and *act>Cas9.P* were completely and partially lethal, respectively (Figure S1C). We reasoned that the very high *Cas9* expression levels that result from expressing UAS-Cas9 in combination with a strong PG-specific Gal4 drivers causes substantial cytotoxicity. Since the PG is responsible for producing ecdysteroids, high levels of Cas9 may interfere with ecdysteroid production and thus disrupt larval and pupal development. Similarly, PG-specific expression of *dCas9.VPR* had only 45% surviving adults, indicating that the toxicity is not necessarily linked to chromosomal breaks, as dCas9 does not cut DNA (Figure 2A). Taken together, these data indicate that combining PG-specific Gal4 with *UAS-Cas9* is not an optimal approach to carry out conditional CRISPR in this tissue, and that other tissues may pose similar issues.

**Figure 2.**
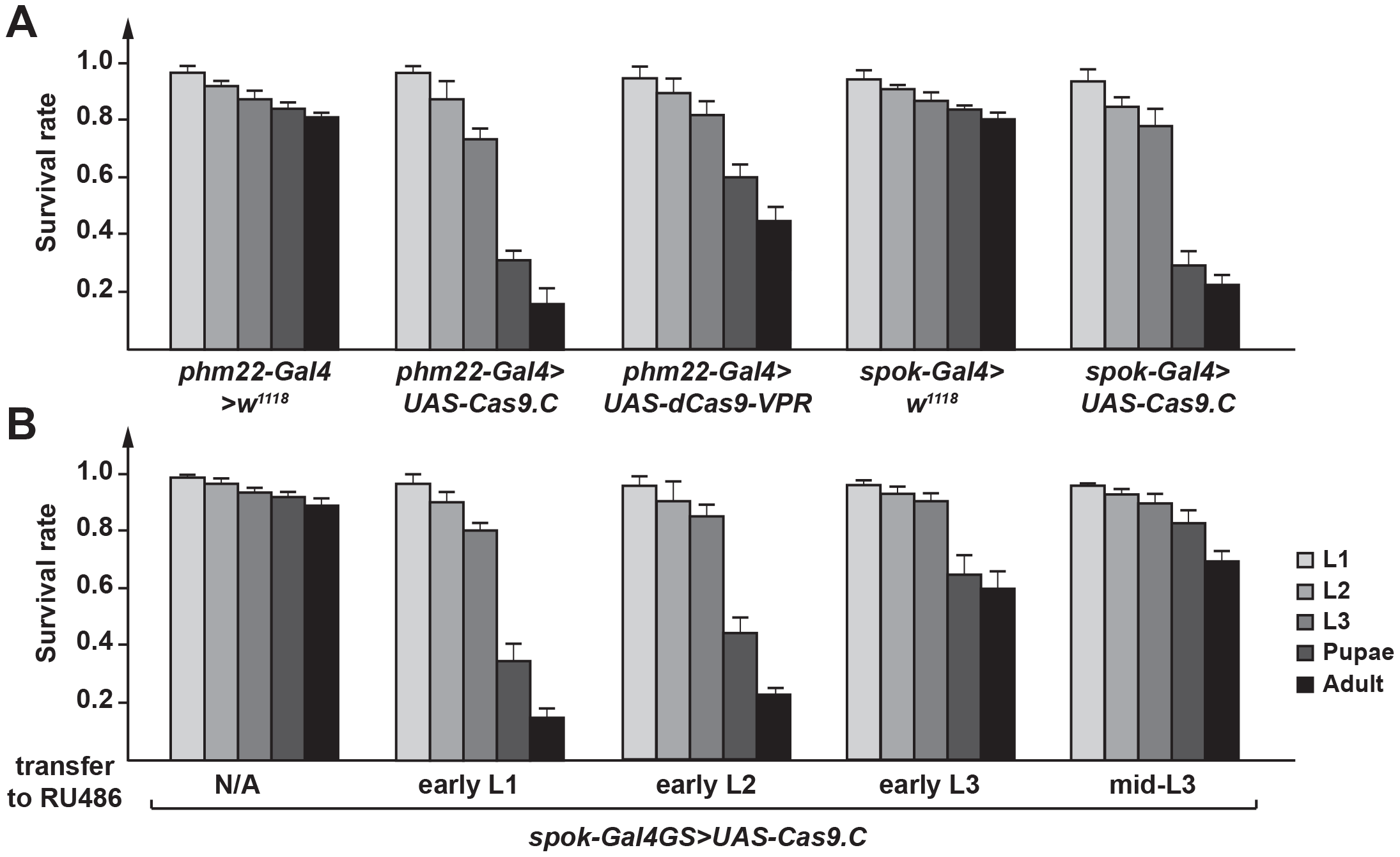
PG-specific Gal4-driven expression of *Cas9* causes lethality. **A.** The survival rates of flies harboring a single copy of *UAS-Cas9* or *UAS-Cas9-VPR* in combination with a single copy of a PG-specific *Gal4* driver (*phm22>* or *spok>*). Error bars represent standard error. The Cas9 cDNA used here is the original allele that is not codon-optimized. UAS-dCas9-VPR is a transgene encoding nuclease-dead Cas9 (dCas9) that is fused to a chimeric co-activator domain (comprising VP64, p65 and Rta). **B.** The survival rates of flies harboring a single copy of *UAS-Cas9* in combination with a single copy of a PG-specific *Gal4GS* (Gal4-GeneSwitch) driver. GAL4GS was activated at different developmental time points by transferring larvae to RU486-supplemented media. Survival rates were quantified for each larval stage and represent surviving animals relative to the number of embryos used per condition (50 embryos for each replicate, three replicates in total). Error bars represent standard error.

**Figure 3.**
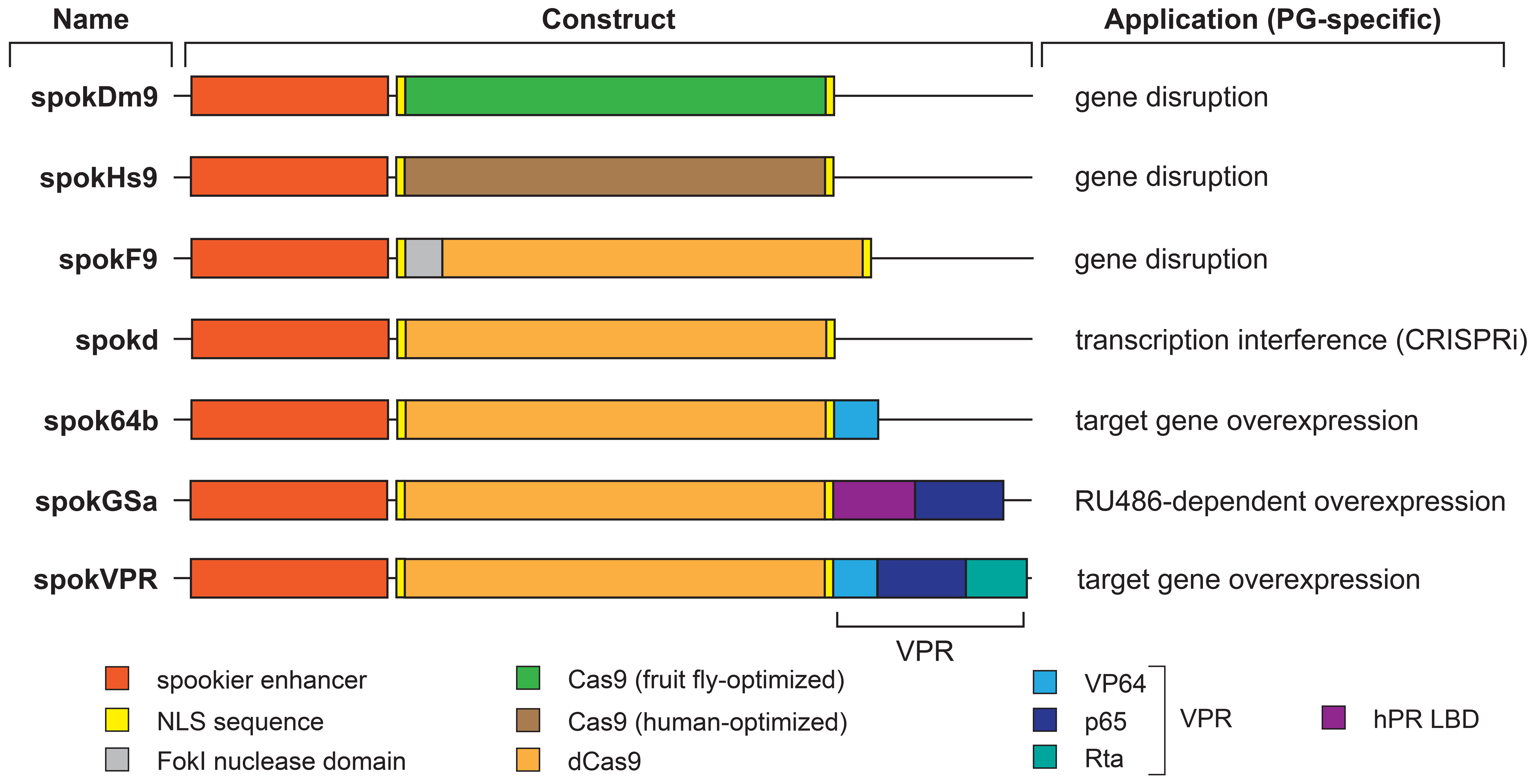
The PG-specific Cas9 (PG-Cas9) vector collection. All vectors are based on the general Gateway Cas9 vector collection (Figure S2). Each PG-Cas9 vector backbone (not shown) is composed of a mini-*white* gene as a marker, a *PhiC31* integrase-compatible *attB* site, the *bla* coding sequence to mediate ampicillin resistance, and a synthetic core promoter. Shown here are the *spookier* (*spok*) regulatory region, the Cas9 variant, the regions encoding Nuclear Localization Sequences (NLS), activations domains (VP64, p65 and Rta), the human Progesterone Receptor Ligand-Binding Domain (hPR LBD) and the FokI nuclease domain. spokDm9, spokHs9 and spokF9 can be used to generate somatic mutations. spokDm9 uses a fruit fly codon-optimized Cas9 version, while spokHs9 is optimized for human cells. spokF9 cuts DNA upon FokI-mediated dimerization followed by FokI cleavage, since dCas (= dead Cas9) is unable to cut DNA. However, the spokd vector harbors dCas9 and can be used to interfere with transcription (CRISPRi) by guiding Cas9 into the vicinity of transcriptional start sites where it may block the assembly of the pre-initiation complex. spok64b, spokGSa and spokVPR were designed to achieve upregulation of target genes. *spokGSa* (GeneSwitch activation) encodes a protein where Cas9 is fused to the hPR LBD and p65 domain, allowing activation via RU486.

#### b. GeneSwitch Gal4/UAS-Cas9

To bypass the toxicity associated with high levels of Cas9, we tested whether temporally controlling Gal4 via the GeneSwitch (GS) system would resolve the problem. The GS system is based on a Gal4 DNA-binding domain that is fused to the human progesterone receptor ligand-binding domain and the activator domain from human p65 (Nicholson et al., 2008). The chimeric Gal4 protein is only activated in the presence of the steroid mifepristone (RU486), which is provided in the diet. Using PG-specific *spok-Gal4GS*, we activated Gal4 during the first (L1), second (L2), third (L3) or mid-third instar larval stages by transferring larvae to a RU486-supplemented diet. Temporal activation of Cas9 as late as early second instar caused still substantial lethality, while later stages (early and mid L3) displayed 60-70% survival. This would make it possibly a suitable approach for studying gene function at later stages but would likely not be ideal for most genes expressed throughout larval development, such as the Halloween genes, which encode ecdysteroid-producing enzymes (Figure 2B). However, given that the PG is a polytene tissue, it remains unclear whether inducing Cas9 during the L3 stage can efficiently disrupt gene function (Ashburner and Richards, 1976).

### The pG-Cas9 system to generate condition CRISPR at tissue of interest

We reasoned that omitting Gal4 altogether and instead opting for endogenous regulatory regions may result in lower but equally specific expression of Cas9, and thus reduce its toxicity. To accomplish this, we sought to generate a vector that would allow for quick insertion of tissue-specific enhancers. For this, we used the existing pBPGUw plasmid, a modular gateway-compatible Gal4 vector (Pfeiffer et al., 2008; Pfeiffer et al., 2010) and replaced the Gal4 sequence by a fragment encoding Cas9 or variants thereof (Figure S2). In brief, DmCas9, HsCas9 and FokI.dCas9 function by generating double-strand breaks and deletions, while dCas9 acts via transcriptional interference. On the other hand, dCas9.VP64, dCas9.GSa and dCas9.VPR are designed to overexpress target genes (Figure S2). More specifically, DmCas9 is codon-optimized for Drosophila, while HsCas9 is codon-optimized for humans (and identical to the afore-mentioned Cas9.P and Cas9.P2, respectively). FokI.dCas9 is a fusion of dCas9 with the nuclease domain of FokI and designed to cut target DNA upon dimerization of the FokI nuclease domain (requires two gRNAs ~ 15-25 bp apart). dCas9 is designed for transcription interference (CRISPRi), where promoter-bound dCas9 will not cut DNA but rather sterically inhibit the proper formation of the pre-initiation complex. Taken together, this set of modified Gateway plasmids can be easily adapted to generate specific enhancer/Cas9 combinations, followed by PhiC31-mediated locus-specific transformation. In order to examine the efficiency of this system, we generated PG-specific versions of these vectors.

### Models and survival rate of transgenic lines carry PG-specific expression of Cas9

For each of the above-listed vectors, we generated corresponding versions that express Cas9 and its variants under the control of the *spookier* (*spok*) regulatory region, which mediates highly specific expression in the PG (Figure 4A) (Ono et al., 2006; Sztal et al., 2007). When we examined the survival rates of these transgenic lines, we noticed that populations heterozygous for any of the *spok-Cas9* constructs were healthy and showed no significant adulthood lethality compared to controls (Figures 2, 4B-C). However, homozygous animals, with the exception of *spokGSa*, displayed some lethality during larval development and substantial or complete lethality during late larval and early pupal stages (Figure 4B-C). In conclusion, heterozygous transgenic lines are viable and can be kept as balanced stocks. We will discuss the efficiency of each construct separately in later sections.

**Figure 4.**
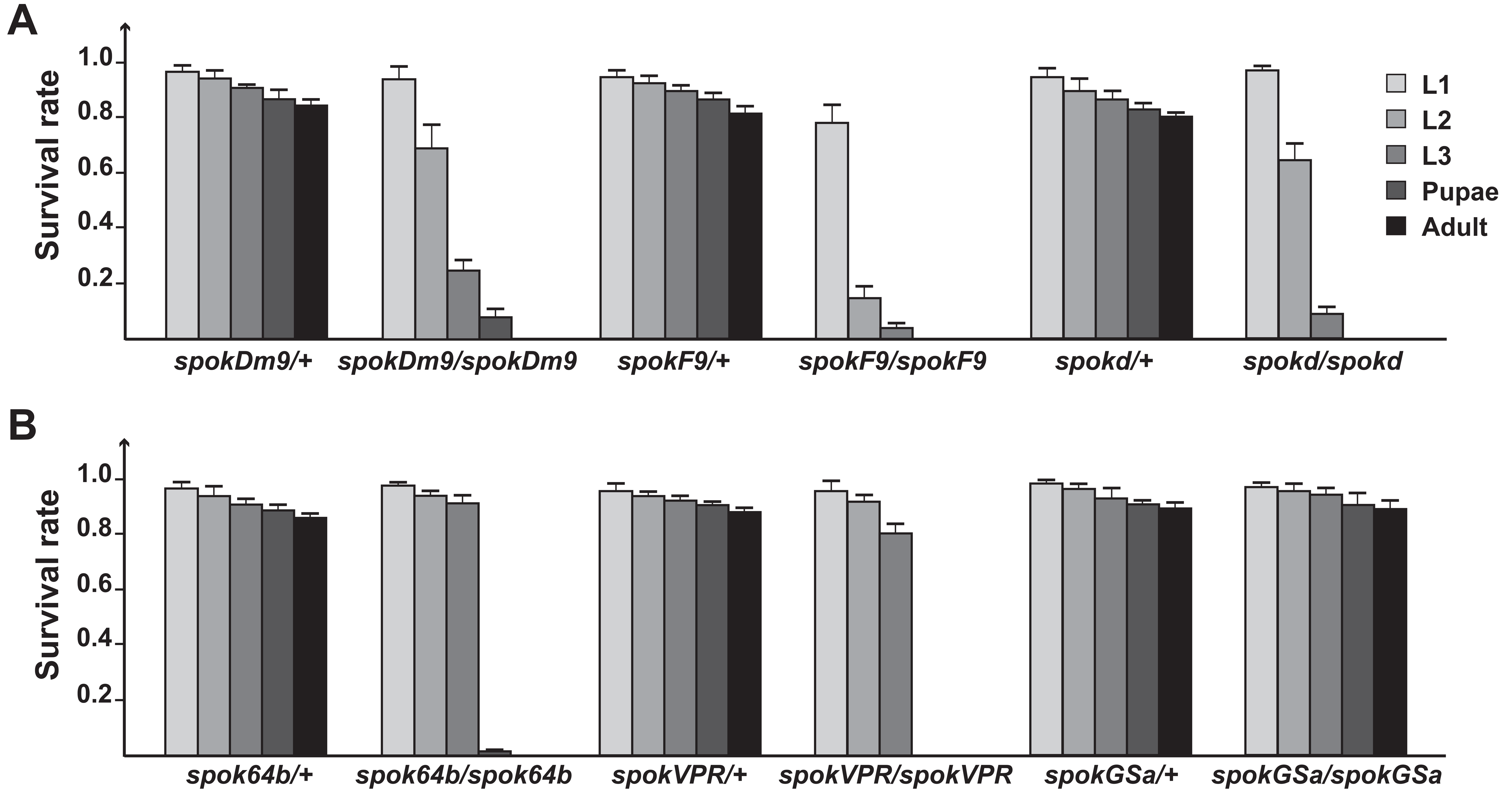
PG-specific expression of Cas9 without Gal4. **A.** The survival rates of flies harboring a single copy or two copies of *spokDm9, SpokF9 or spokd* or *UAS-Cas9-VPR*. Error bars represent standard error. **B.** The survival rates of flies harboring a single copy or two copies of *spok64b, SpokVPR or spokGSa*. Error bars represent standard error. For A and B data was normalized to the number of embryos in the starting population.

### Localization of Cas9 in the PG

Before examining whether our Cas9 transgenes caused PG-specific alterations in gene expression, we first examined the presence of Cas9 protein in PG nuclei. Previous studies have shown that epitope tags might affect the DNA-binding properties of Cas9, which prompted us to remove the 3xFlag tags found in the original dCas9.VPR plasmid, which ensured that all Cas9 transgenes were untagged. As expected, immunostaining with anti-Cas9 antibodies showed robust presence in PG nuclei, while the expression in nearby tissues, including the CA and CC was negligible (Figure 5).

**Figure 5.**
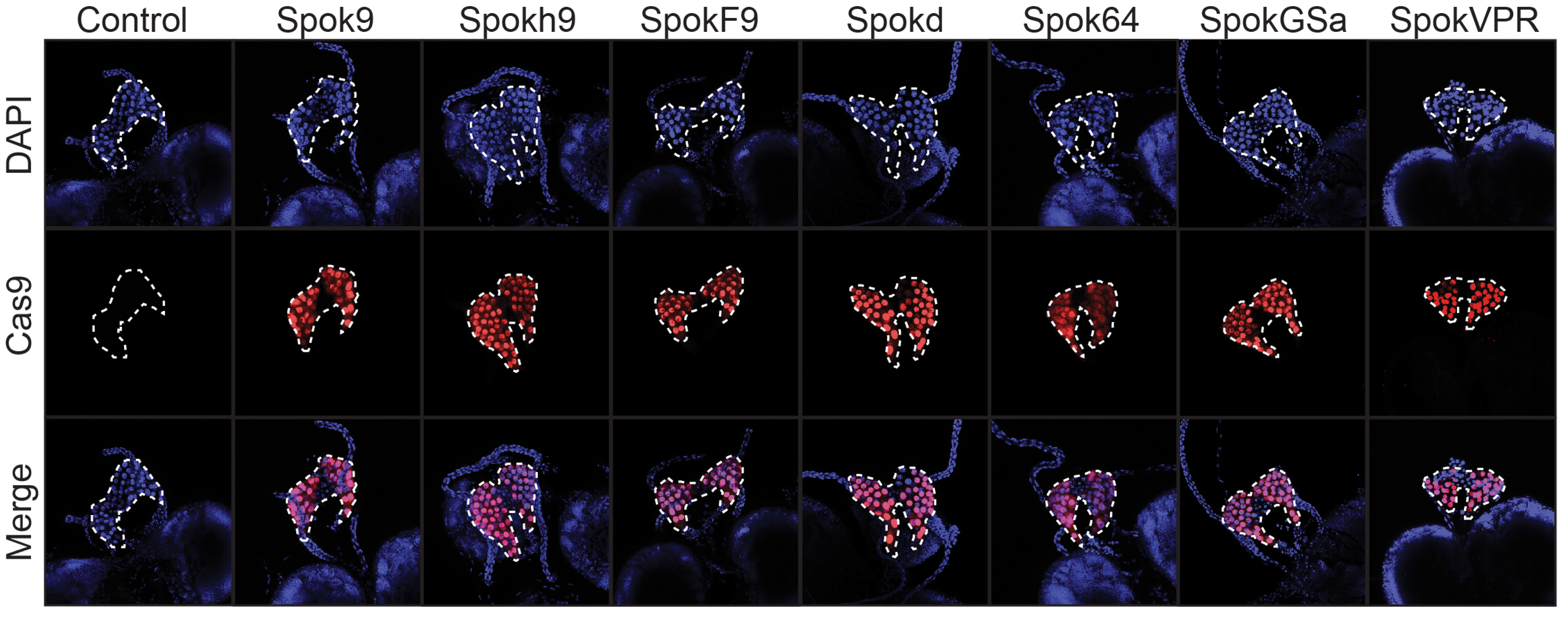
Nuclear localization of Cas9 in the *Drosophila* prothoracic gland (PG). Nuclear localization sequences were added to the 5’ and 3’ ends of the *Cas9* cDNA to ensure transport of Cas9 into nuclei. The *spok* regulatory region drives the expression of Cas9 specifically in PG cells with no detectable signal in the adjacent corpora allata and the corpora cardiaca. DAPI (blue) was used to stain DNA while anti-Cas9 antibodies (red) was used to detect Cas9.

### Mutation efficiency of PG-specific gene disruption via Cas9

The ability to generate somatic gene mutations is still limited in *Drosophila* and has not been reported for the PG. We therefore used three different strategies to generate transgenes with PG-specific Cas9 expression, comprising spokDm9 (fly codon-optimized), spokHs9 (human codon-optimized) and spokF9 (FokI nuclease domain fused to fly codon-optimized dCas9). In order to test the efficiency for each approach, we generated transgenic lines carrying gRNAs against either *phantom* (*phm*) or *disembodied* (*dib*), two well characterized genes involved in ecdysteroid synthesis (Niwa et al., 2004; Warren et al., 2004). Classic mutants of *phm* and *dib* display embryonic lethality, while PG-specific *phm*- and *dib*-RNAi cause L1 and L3 arrest, respectively. Both *phm*- and *dib*-RNAi populations can be rescued to adulthood when reared on 20E-supplemented media (Figure 6A-B) (Niwa et al., 2004; Niwa and Niwa, 2016), and we reasoned that the specificity of *phm*- and *dib*-gRNAs could be easily assessed by 20E-feeding as well. To generate double-strand breaks (DSB) in the coding region of *phm* or *dib*, we generated transgenic lines that carried at least two gRNAs (*dib*^*gR1*^ and *phm*^*gR1*^), where the distance of the target sequence would not exceed 400 bp (Table S1, Figure S3). For spokF9, DSBs are not achieved by the endonuclease activity of Cas9 (which is missing in dCas9) but require dimerization of the FokI nuclease domain, a bacterial type IIs restriction enzyme (Wah et al., 1997). Dimerization of FokI is dependent on the recruitment of two Cas9 molecules guided by two distinct gRNAs that are 15-25 bp apart. We therefore generated transgenic lines that carry two pairs of gRNAs to allow for Cas9-FokI-mediated deletions (Table S1, Figure S3).

**Figure 6.**
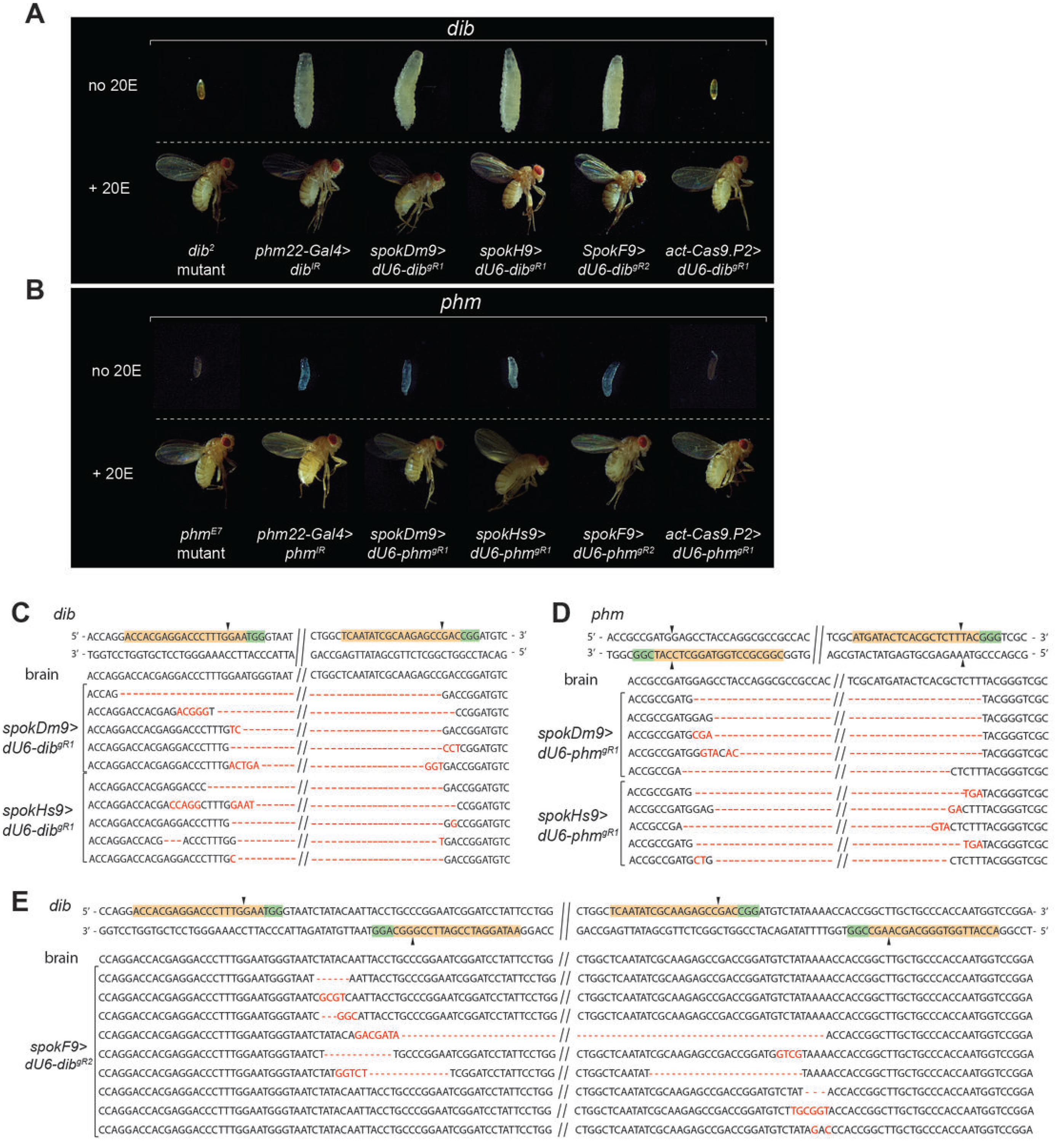
Efficiency of tissue-specific CRISPR/CAS9 in the *Drosophila* prothoracic gland (PG). **A.** Comparing phenotypes of a classic *disembodied* mutant (*dib*^*2*^) and PG-specific RNAi (*dib*^*IR*^) with PG-specific (*spokDm9, spokHs9* and *spokF9*) or ubiquitous CRISPR/Cas9 (*act-Cas9.P2*). **B.** Comparing phenotypes of a classic *phantom* mutant (*phm*^*E7*^) and PG-specific RNAi (*phm^IR^*) with PG-specific (*spokDm9, spokHs9* and *spokF9*) or ubiquitous CRISPR/Cas9 (*act-Cas9.P2*). **C and D.** Sequences of *dib* (C) and *phm* (D) loci from brain and PG nuclei, using either *spokDm9* or *spokHs9*. **E.** Sequences of *dib* locus from brain and PG nuclei using *spokF9* in combination with *dU6-dib^gR2^*

In summary, using either spokDm9, spokHs9 or spokF9 to induce PG-specific DSBs in the *phm* and *dib* genes yielded similar results, and all caused phenotypes that were similar to those seen in *phm*- and *dib*-RNAi animals. The Cas9 lines were less leaky than the RNAi approach, with very few pupal and no adult escapers (Table S2). Importantly, Cas9/gRNA animals were rescued to adulthood when reared on a diet supplemented with 20E, with typically 70-80% of the population developing into adults (Table S2), indicating that the phenotypes resulted specifically from gene disruptions in *phm* and *dib* (Figure 6A-B).

To further confirm the specificity of the *dib* and *phm* gRNAs, we crossed either line to *act-Cas9*, allowing us to target *dib* and *phm* in a ubiquitous manner, which should give rise to phenotypes that are similar to the corresponding classic mutants. In agreement with this, both *act-Cas9.P2>phm*^*gR1*^ and *act-Cas9.P2>dib*^*gR2*^ were embryonic lethal, and thus phenocopied the classic mutants (Figure 6A-B). To ensure that these phenotypes were indeed caused by a disruption of the intended target genes, we extracted genomic DNA from hand-dissected ring glands and sequenced the *phm* and *dib* gene regions. As a control, we isolated genomic DNA from the adjacent brain. Upon sequencing at least 10 clones per line we found that both Drosophila-and human-optimized Cas9 (spokDm9 and spokHs9) in combination with two gRNAs were highly efficient in generating deletions in the predicted region (Figure 6C-D). Some of clones appeared to be wild type alleles (not more than three out of ≥10 per line, not shown), however, since the ring gland samples comprised two non-targeted tissues (the corpora cardiaca and the corpora allata, Figure 1), we cannot distinguish between loci that were not targeted in the PG and loci that originate from the other two Cas9-free cell types. In comparison to spokDm9 and spokHs9, using spokF9 in combination with two gRNA pairs resulted in fewer large deletions, suggesting this approach was less efficient in this regard. However, on a phenotypic level, spokF9 was just as efficient as spokDm9 and spokHs9, all of which were 100% lethal (Table S2). All tested clones derived from brain samples were wild type, indicating that the *spok* regulatory region does promote little or no expression in brain cells.

### *In vivo* transcription interference via PG specific dCas9 (spokd)

CRISPR applications are not limited to ablating gene function via DSBs. An alternative strategy is to interfere with the transcription of a target gene (CRISPRi). This is a desirable approach, since RNAi may either not efficiently repress gene function or result in off-target effects (Jackson and Linsley, 2010; Bier et al., 2018). CRISPRi utilizes gRNAs that will recruit endonuclease-dead Cas9 (dCas9) to promoter regions of target genes, ideally close to the transcription start site (TSS), which will interfere with the assembly of the transcription machinery at this locus (Larson et al., 2013; Ghosh et al., 2016). This strategy is ideal for selectively targeting specific promoters of genes that harbor alternative promoters to repress specific mRNA isoforms. Alternatively, one could interfere with the expression of a gene for a defined duration, and then revert back to normal expression, thus studying dynamically expressed genes. We wanted to test whether PG-specific CRISPRi would work as efficiently as the other tools at our disposal. As candidate genes, we again chose *phm* and *dib*. We generated lines expressing a single gRNA targeting either −423 or −174 bp upstream of the *phm* TSS, and for *dib* we selected −482 and −110 bp upstream (Figure S3). When we crossed these four gRNA lines to flies carrying *spokd* transgenes, we observed developmental arrest during the L3 stage, which, in the case of *dib*, was comparable to what we had observed in *phm22-Gal4>dib*-RNAi animals (Figure 7A-B) and the corresponding Cas9-driven gene knockouts (Figure 6A). In contrast, targeting *phm* via CRISPRi, while lethal, was less efficient compared to the other strategies, as larvae died at later stages. This may suggest that the chosen gRNA sites were too far away from the *phm* TSS, since no alternative promoters have been reported for this gene. However, based on qPCR analysis, the relative reduction of transcript levels in the CRISPRi lines were comparable between *phm* and *dib*, as we observed a 2- to 6-fold reduction for *phm* and a 2.5- to 4-fold reduction for *dib* (Figure 7B). It is possible that *phm* transcript levels need to even stronger reduction to elicit phenotypes that are comparable to the Cas9 knockouts. Finally, to ensure that these phenotypes arise not from DSBs, we sequenced these loci and found them to be wild type in all cases (Figure 7C).

**Figure 7.**
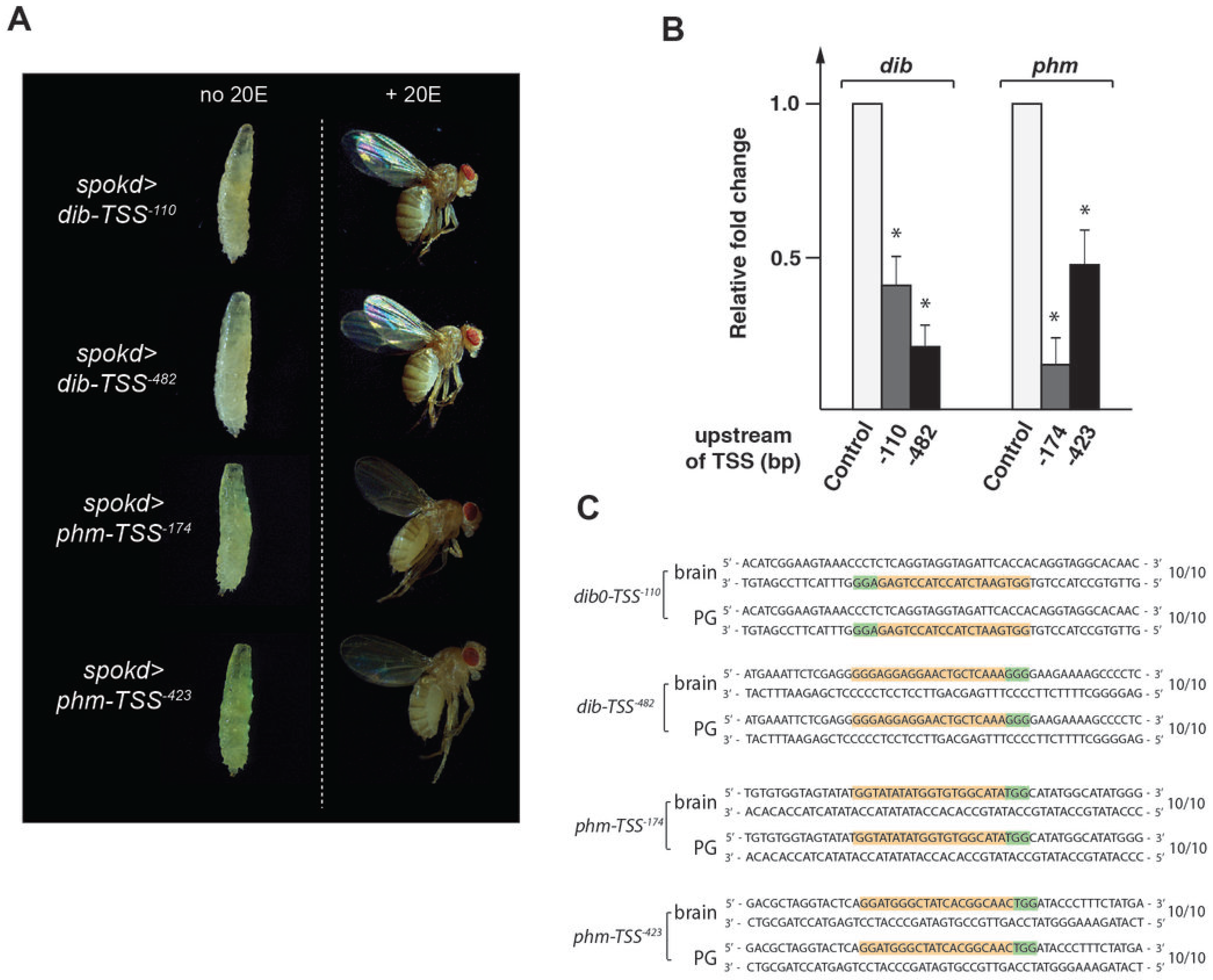
Efficiency of PG-specific CRISPR interference (CRISPRi) in *Drosophila*. **A.** *spokd* (= *spok-dCas9*) is used to ubiquitously express gRNAs targeting −110 and −482 bp upstream of *dib* as well as −174 and −423 bp upstream of *phm* (relative to TSS), respectively. In each case, this resulted in L3 arrest (left) and rescue to adulthood when the diet was supplemented with 20E (right). For comparison to PG-specific *dib*- and *phm*-RNAi and classic mutant phenotypes, see Figures 6A-B. **B.** RG-specific qPCR for *dib*- and *phm*-CRISPRi. Ring glands were dissected at 42 hrs after the L2/L3 molt, three replicates per condition. * => p-value < 0.05. **C.** Sequences of *dib* and *phm* loci obtained from DNA of CRISPRi-treated PG nuclei show no alterations. For each condition, we sequenced 10 clones, all of which were wild type.

### Upregulating gene expression via PG-specific CRISPR/Cas9

Previous approaches aimed at overexpressing a specific gene were based on the generation of transgenic lines that carry a cDNA, either driven by heat-shock promoters, nearby enhancers or the Gal4-UAS system (Pirrotta, 1988; Stapleton et al., 2002; Busson and Pret, 2007). Later improvements included the use of the PhiC31 system to ensure locus-specific integration and consistent expression of the transgene (Fish et al., 2007). However, these approaches require the generation and cloning of a cDNA, which may be time-consuming and difficult. Using dCas9 variants that harbor activation domains, one can now direct dCas9 to specific endogenous promoters and activate any given target gene, referred to as CRISPRa (a= activation). We therefore generated PG-specific versions of dCas9, to which we fused the VP64 or VPR activation domains (Lin et al., 2015; Dominguez et al., 2016), named here spok64b and spokVPR (short for spokdCas9-VP64b and spokdCas9-VPR) (Figure 4A). A report by the Perrimon lab showed that dCas9-VP64 was not as efficient as dCas9-VPR to activate target genes (Lin et al., 2015). However, the dCas9-VP64 construct only contained two nuclease-attenuating mutations D10A and H840A compared to the dCas9-VPR, which contained four (D10A, H839A, H840A and N863A). We therefore modified the original *dCas9-VP64* to *dCas9-VP64b*, so that it contained the same four nuclease-attenuating mutations as the *dCas9-VPR* construct.

We first examined the efficiency of the *spok64*, *spok64b* and *spokVPR* constructs by transfecting cultured brain-ring gland complexes (BRGC) that carried gRNA transgenes targeting either the *phm* or *dib* promoters upstream of the TSS, which were the same lines as used for CRISPRi. Since the plasmid-encoded *Cas9* alleles were driven by the *spok* regulatory region, we reasoned that this approach should result in PG-specific *Cas9* expression. Indeed, when we used ring gland-specific qPCR, *spokVPR* resulted in a 10- to 30-fold induction, while *spok64b* ranged from 5- to 15-fold upregulation. In contrast, the *spok64* plasmid showed essentially no increased gene expression, suggestion that the two additional point mutations in *spok64b* (839A and N863A) are critical for induction (Figure 8A). In order to ensure that *spok64b* and *spokVPR* worked similarly efficient *in vivo*, we generated corresponding transgenic lines, and crossed them to transgenic lines carrying gRNAs that target regions upstream of the *phm* and *dib* TSS. Similar to our BGRC transfection results, *spokVPR* resulted in a 9- to 28-fold induction, while *spok64b* upregulated expression ranging from 4-to 18-fold (Figure 8B).

**Figure 8.**
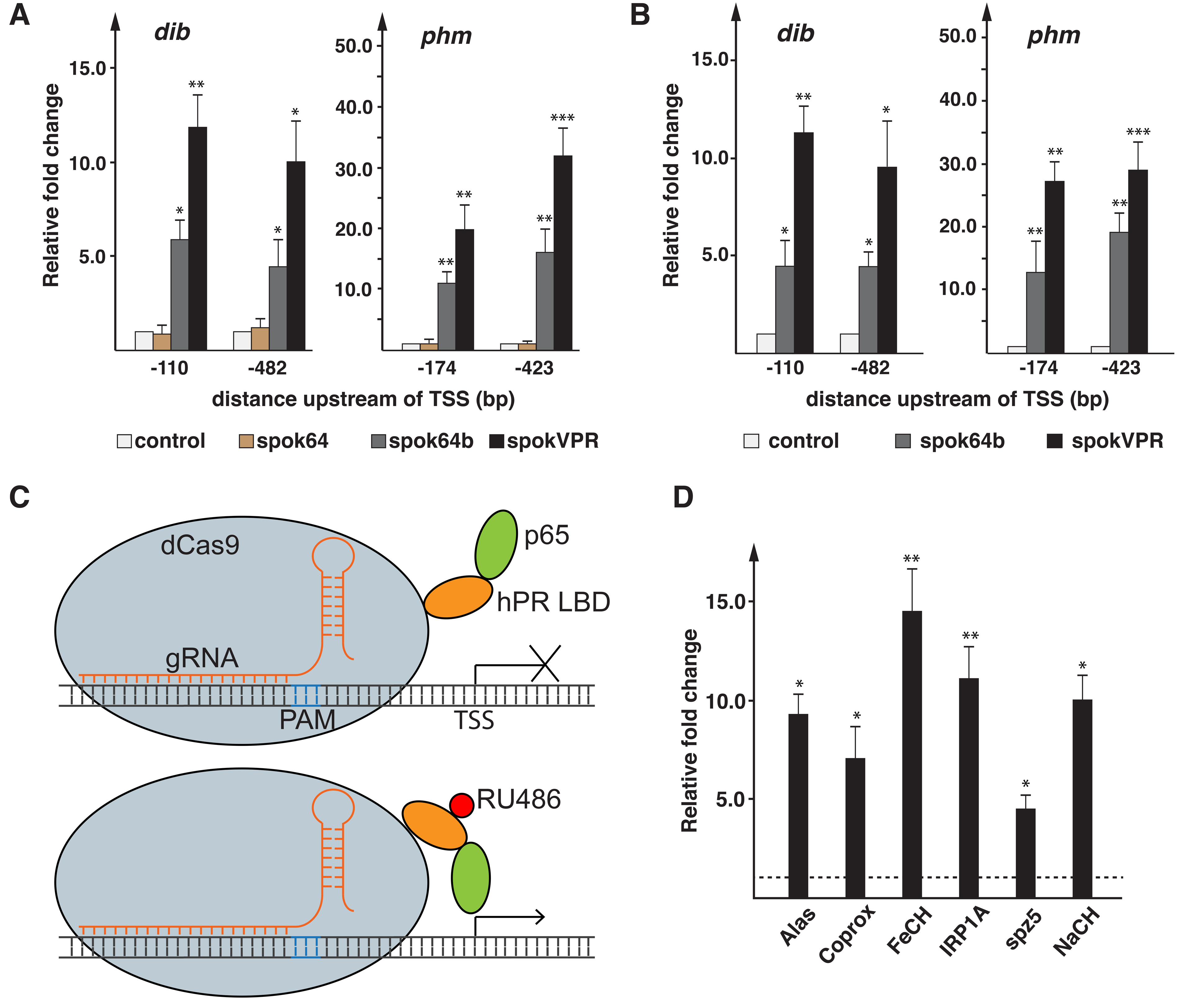
Efficiency of PG-specific CRISPRa. **A.** qPCR of *ex vivo*-cultured ring glands transfected with *spok64, spok64b* and *spokVPR* plasmids. Transfected glands were isolated from transgenic larvae expressing gRNAs targeting −110 and −482 bp upstream of *dib* as well as −174 and −423 bp upstream of*phm* (relative to TSS), respectively. All results normalized to controls (= no plasmid added). spok64 differs from spok64b, as it encodes different amino acids at positions 839 and 863, which are important for attenuation of the endonuclease, as previously reported (Lin et al., 2015). *spokVPR* has the same changes as *spok64b*, but in addition harbors p65 and Rta domains (Figure 3). **B.** Same as A, however ring glands were transgenic for both gRNA and Cas9 constructs. *spok64* was not used to make transgenics, due to the lack of activity shown in A. **C.** Schematic of dCas9 fused to the human progesterone ligand-binding domain (hPR LBD) and the p65 activation domain, resulting in dCas9GSa (= dead Cas9-GeneSwitch for activation). This approach allows for temporal control over the activation via dCas9, by switching animals to a diet supplemented with RU486. **D.** qPCR analysis of six target genes. Lines were obtained from Bloomington stock centre. Shown are the fold-changes relative to the same gene in samples of the same genotype, but raised on RU-486-free medium (dotted line = 1). Ring glands were dissected from larvae that were reared for four hours on media ± RU486. * = p-value < 0.05, ** = p-value < 0.01, *** = p-value < 0.001. Error bars represent standard error.

Finally, we also generated a version of CRISPRa that allows for temporally-controlled gene induction. We wondered whether we could render Cas9 ligand-inducible, similar to the GeneSwitch (GS) system, where the Gal4 DNA-binding domain is fused to the human progesterone receptor-ligand-binding domain (hPR-LBD) and the p65 activation domain. The resulting chimeric Gal4 protein can only be activated in the presence of steroid mifepristone (RU486), typically provided in the diet. Therefore, we cloned a *spok*-driven chimeric cDNA encoding the catalytically inactive dCas9 fused to hPR-LBD and p65 (Figure 8C) and generated the corresponding transgenic line (aka spokGSa). In order to assess the efficiency of RU486- mediated induction, we chose target genes we are actively studying in the lab (*Alas*, *Coprox*, *FeCH*, *IRP1A*, *spz5*, *Nach*) and that have comparatively flat expression profiles in the PG compared to *phm* and *dib* during larval development (Ou et al., 2016). When we crossed the corresponding gRNA transgenic lines to *spokGSa* and switched larvae to a RU486-containing diet, we observed PG-specific upregulation as early as two hours after exposure to RU486, similar to what has been reported in Gal4GS system (Figure 8D) (Nicholson et al., 2008). After four hours, induction of target genes ranged from 4- to 15-fold compared to controls, indicating that the GeneSwitch system works well, and is a powerful tool to temporally control gene upregulation.

### Using PG-specific gRNAs for modulating gene expression

To manipulate gene expression in a tissue-specific manner via CRISPR, one can utilize two main strategies: (i) restricting Cas9 expression to specific tissues or (ii) limiting the expression of gRNA to the tissue of interest. In the approaches outlined above, we employed PG-specific Cas9 expression. We therefore tested whether reversing gRNA and Cas9 expression patterns from tissue-specific to ubiquitous (and vice versa) was a viable strategy, because an existing line with ubiquitous Cas9 expression was reported to be homozygous viable (Bloomington stock #58590). A previous study described using UAS-driven multiplexed gRNA cloned into pCFD6 (Addgene 73915) to mediate tissue-specific gRNA expression in *Drosophila* imaginal wing discs (Port and Bullock, 2016). In contrast to pCFD6, other gRNA-generating vectors use U6-type promoters (pU6), which are RNA Polymerase III promoters that drive ubiquitous expression of gRNAs (Port et al., 2014; Port and Bullock, 2016). This pU6-based approach has the potential to cause non-specific mutagenesis in non-target tissues where Cas9 expression is leaky (Port and Bullock, 2016). However, since pCFD6 requires an additional Gal4-expressing transgene, and building the corresponding fly lines to achieve tissue-specific lesions is thus more complex.

In an effort to improve available tools for tissue-specific gRNA production, we replaced the pU6:3 promoter in the commonly used pCFD5 plasmid (Port and Bullock, 2016) with the *spok* regulatory region and added sequences mediating hammerhead (HH) or Hepatitis delta virus (HDV) ribozyme function to induce self-cleavage and proper release of gRNAs (Webb and Lupták, 2011; Manivannan et al., 2015) (Figure 9A). The pCFD5 plasmid harbors two Gly-tRNA sequences that allow insertion of two gRNAs. However, additional tRNA-gRNA pairs can be added if one requires more than two gRNAs, which we recommend for targeting large genes or if one wishes to target multiple genes with a single construct. Unlike pU6 promoters, the *spok* enhancer recruits Polymerase II (Pol II), resulting in mRNAs that will be subjected to 5’-capping and 3’-polyadenylation, which have the potential to interfere with proper gRNA maturation (Arimbasseri et al., 2013; Darnell, 2013). Therefore, we added the HH and/or HDV ribozyme sequences to three of our vectors (PG2-4) to test whether this would result in more efficient phenotypes due to increased processing of gRNAs, while one vector (PG1) received no ribozyme sequence (Figure 9A). As a consequence of this design, the resulting transgenic lines require only a single cross to combine Cas9 and gRNA in the F1 generation. The use of ribozyme sequences has been successfully used in zebrafish and *Arabidopsis* (Lee et al., 2016; Zhang et al., 2017), but has not been described in *Drosophila*. Compared to the original pCFD5 vector, all cloning steps for our PG-gRNA plasmids are exactly the same, requiring no additional adjustments in terms of cloning strategy (Port and Bullock, 2016).

**Figure 9.**
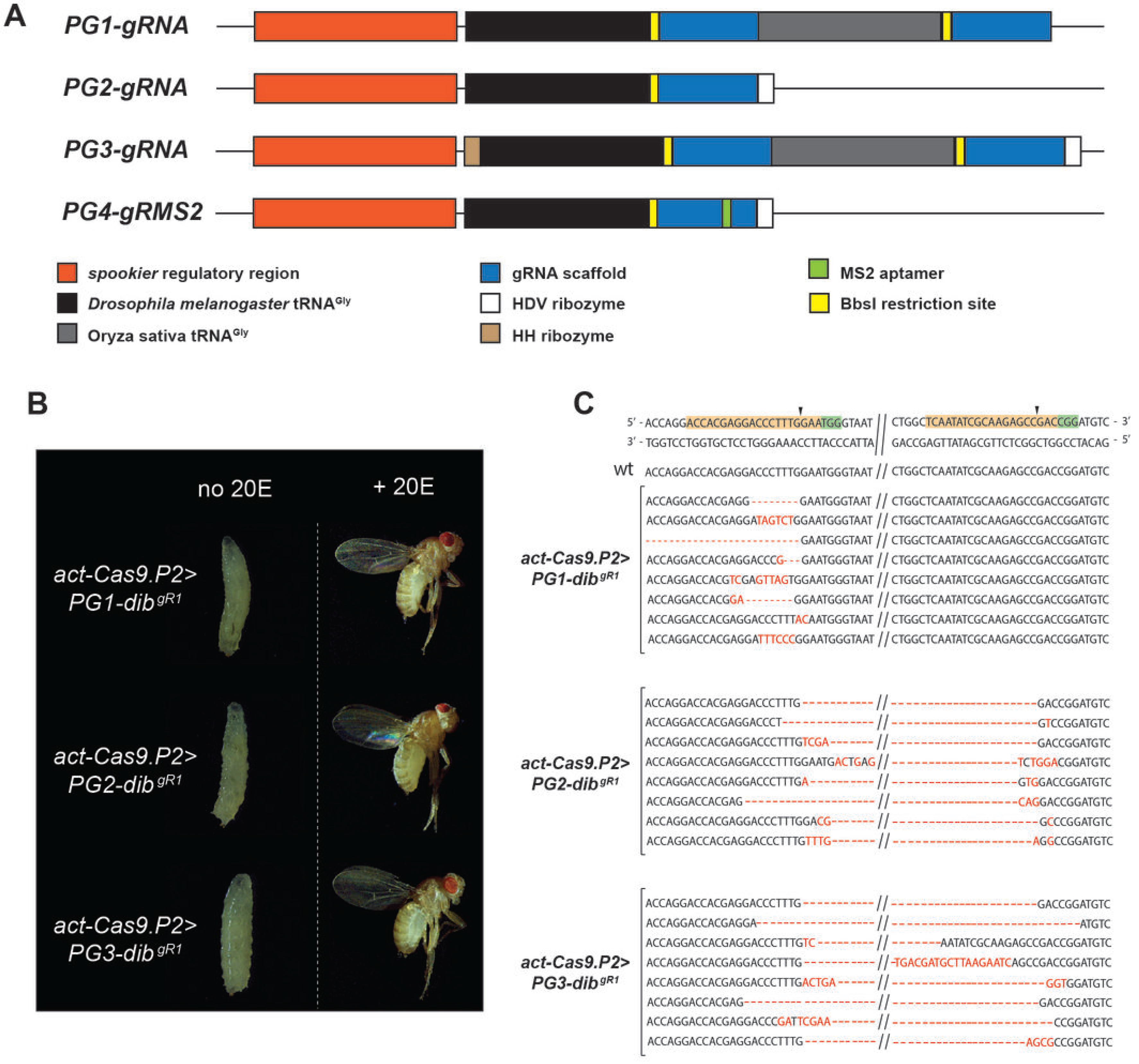
PG-specific gRNA in combination with ubiquitous Cas9 expression. **A.** Schematic illustration of PG-gRNA constructs that allow for prothoracic gland-specific expression of multiple gRNAs in a single vector, which are based on the commonly used pCFD5 plasmid where we replaced the pU6:3 promoter with the *spok* regulatory region. We also added sequences mediating hammerhead (HH) or Hepatitis delta virus (HDV) ribozyme function to promote proper processing of gRNAs from Pol II-derived mRNAs (Webb and Luptak, 2011; Lee et al., 2016). Like the original pCFD5 plasmid, this vector series harbors two tRNA^Gly^ sequences that natively allow the insertion of two gRNAs, but additional gRNA-tRNA fragments can be added to target larger regions of DNA. PG1-3 are used for gene disruption, while PG4 (not tested in this study), which harbors an MS2 aptamer, is intended for gene activation in combination with fly lines that carry MCP_p65_hsf1 or MCP_VP64 transgenes (“flySAM”). **B.** Phenotypes associated with PG-specific gRNA-and ubiquitous Cas9-expression targeting the *dib* gene, in the absence (left) or presence (right) of dietary 200H-ecdysone (20E). **C.** Sequences of *dib* locus resulting from using the same *dib* gRNA pair (gR1), but different PG-gRNA vectors (PG1-3). Red letters and dashes indicate altered or missing nucleotides.

In total, we generated four different PG-gRNA vectors, three of which are designed for generating DSBs (PG1-3), while one them (PG4) harbors an MS2 aptamer (Chavez et al., 2016) to mediate gene upregulation (Figure 9A). For PG1-gRNA, we did not add any ribozyme sequences, while PG2-gRNA has an HDV ribozyme sequence at the 3’ end and PG3-gRNA harbors an HH ribozyme at the 5’ end as well as HDV region at the 3’ end of the multiplex. In order to evaluate the efficiency of this approach, we inserted the same two gRNAs targeting the *dib* gene that we used for ubiquitous pCFD5-driven gRNA expression, ensuring that any differences in phenotypes arise from ubiquitous vs. tissue-specific gRNA expression (Table S1, S2, Figure S3).

When we crossed transgenic lines carrying either *PG1-dib*^*gR1*^, *PG2-dib*^*gR1*^ or *PG3-dib*^*gR1*^ to Act-Cas9, we observed consistently 100% L3 arrest, similar to *spok9>pU6-dib*^*R1*^ animals that produced ubiquitous gRNA and PG-specific Cas9. In addition, supplementation with 20E efficiently rescued the L3 lethality, resulting in 72-82% normal-looking adults, indicating the specific disruption of the *dib* gene (Figure 9B and Table S2). However, when we sequenced genomic DNA from dissected ring glands, we noticed that the *PG1-dib^gR1^* transgene, which lacked the ribozyme sequences, was less efficient compared to *PG2-dib^gR1^* and *PG3-dib^gR1^*. Specifically, while *PG2-dib^gR1^* or *PG3-dib^R1^* consistently caused deletions resulting from mutations at both gRNA loci, the *PG1-dib^gR1^* construct failed to produce mutations for the downstream gRNA and therefore lacked corresponding deletions (Figure 9C). This suggests that the addition of the 3’ HDV ribozyme sequences was necessary to allow for effective processing of the downstream gRNA. Taken together, these data demonstrated that restricting the expression of gRNA to the PG is highly effective and provides an alternative solution to the lethality issue caused by PG-specific Cas9 expression, provided one uses ribozyme sequences to ensure appropriate processing of gRNAs from Pol II-derived mRNAs.

We have not tested the functionality of the PG4 construct, which harbors the MS2 aptamer, but make it available for public testing. The addition of the MS2 sequence promotes the recruitment of the MS2 coat protein (MCP) (Chavez et al., 2016) and MCP fusion proteins such as MCP_p65_hsf1 and MCP_VP64, which are available as transgenic fly lines (“flySAM”) (Jia et al., 2018). The binding of MCP-coactivator fusions is aimed at causing stronger gene upregulation compared to VP64 and VPR alone.

## Conclusions

We showed here that PG-specific expression of Cas9 causes lethality that is independent of its nuclease activity. We have generated a series of strategies to solve this problem, which now allows somatic generation of DSBs, CRISPRi and CRISPRa. Generating tissue-specific gRNAs is also a viable strategy, provided ubiquitously expressed Cas9 levels are sufficiently low to avoid lethality. Since endoreplicating tissues harbor multiple copies of the same locus, Cas9 should be activated early enough to ensure efficient gene disruption. In our hands, somatic gRNAs work even better than RNAi lines, and appear to be highly specific, indicating that polytene tissues pose no issue for somatic CRISPR approaches. Even though we generated tools for PG-specific CRISPR, our tools can be used for any tissue of interest, polytene or not.

## Acknowledgements

We thank Norbert Perrimon, Simon Bullock, Gerald Rubin and Michael O’Connor for sharing the original plasmids used in this study. This work was support by Natural Sciences and Engineering Research Council of Canada (RGPIN-2018-04357).

## Materials and Methods

### *Drosophila* stocks and husbandry

*y*^*1*^*v*^*1*^*P*(nos-PhiC31.NLS;)*X*; *P*(*carryP*)*attP40(II)* (#25709), *y*^*1*^*v*^*1*^*P*(nos-PhiC31/int.NLS)*X*; *P*(carryP)*attP2(III)*(#25710), *phm*^*E7*^/*FM7c* (#2208), *dib*^*2*^/*TM3 Sb*^*1*^ (#2776), *UAS-Cas9.P2* (#58985), *UAS-Cas9.P* (#54594), *UAS-Cas9.P* (#54595), *Act-Cas9* (#58590), *spz5 P*(OE.gRNA)*attP40* (#67547), *Alas P*(OE.gRNA)*attP40* (#68083), *Coprox P*(OE.gRNA)*attP40* (#68124), *FeCH P*(OE.gRNA)*attP40* (#78206), *IRP1A P*(OE.gRNA)*attP40* (#68039), *Nach P*(OE.gRNA)*attP40* (#67562) were obtained from the Bloomington Drosophila Stock Center. *y*^*2*^*cho*^*2*^*v*^*1*^ (TBX-0004), *y*^*2*^*cho*^*2*^*v*^*1*^; *sco/CyO* (TBX-0007), *y*^*2*^*cho*^*2*^*v*^*1*^/*Y*^*hs-hid*^; *Sp/CyO* (TBX-0008), *y*^*2*^*cho*^*2*^*v*^*1*^; *Sp hs-hid/CyO* (TBX-0009), *y*^*2*^*cho*^*2*^*v*^*1*^; *Pr Dr/TM6C, Sb Tb* (TBX-0010) were obtained from the National Institute of Genetics of Japan (NIG).

*UAS-phm*-RNAi (#108359), *UAS-dib*-RNAi (#101117) were obtained from the Vienna *Drosophila* Resource Center.

*Spok.DmCas9/TM3,Ser.GFP* (SpokDm9), *Spok.HsCas9/CyO.GFP* (SpokHs9), *Spok.Fok1-dCas9/CyO.GFP* (SpokF9), *Spok.dCas9-VP64b/TM6B,Hu,Tb* (Spok64b), *Spok.dCas9-GS-p65* (SpokGSa), *Spok.dCas9-VPR/TM6B,Hu,Tb* (SpokVPR), *Spok.dCas9/TM3,Ser.GFP* (Spokd), *y*^*1*^*v*^*1*^;*P*(pCFD3 no target gRNA)*attP40* (pCFD3 control), *y*^*1*^*v*^*1*^;*P*(pCFD5 no target gRNA)*attP40* (pCFD5 control), *y*^*1*^*v*^*1*^;*P*(PG1.gRNA no target gRNA)*attP40* (pPG1.gRNA), *y*^*1*^*v*^*1*^;*P*(PG2.gRNA no target gRNA)*attP40* (pPG2.gRNA), *y*^*1*^*v*^*1*^;*P*(PG3.gRNA no target gRNA)*attP40* (pPG3.gRNA), *y*^*1*^*v*^*1*^;*P*(pCFD5 phm.KO dgRNA)*attP40* (dU6-phm^gR1^), *y*^*1*^*v*^*1*^;*P*(pCFD5 phm gRNA SpokF9) (dU6-phm^gR2^), *y*^*1*^*v*^*1*^;*P*(pCFD5 phm.5TSS −174 gRNA)*attP40* (phm TSS^−174^), *y*^*1*^*v*^*1*^;*P*(pCFD5 phm.5TSS −423 gRNA)*attP40* (phm TSS^−423^), *y*^*1*^*v*^*1*^;*P*(pCFD5 dib.KO dgRNA)*attP40* (dU6-dib^gR1^),*y*^*1*^*v*^*1*^;*P*(pPG1.dib KO dgRNA)*attP40* (pPG1-dib^gR1^), *y*^*1*^*v*^*1*^;*P*(pCFD5 dib gRNA SpokF9) (dU6-dib^gR2^), *y*^*1*^*v*^*1*^;*P*(pPG2.dib KO dgRNA)*attP40* (pPG2-dib^gR1^), *y*^*1*^*v*^*1*^;*P*(pPG3.dib KO dgRNA)*attP40* (pPG-.dib^gR1^),*y*^*1*^*v*^*1*^;*P*(pCFD5 dib.5TSS-110 gRNA)*attP40* (dib TSS^−110^), *y*^*1*^*v*^*1*^;*P*(pCFD5 dib.5TSS −482 gRNA)*attP40* (dib TSS^−482^) were generated by our lab.

*y*^*1*^*v*^*1*^/*SM5,CyO*, *y*^*1*^*w*\**P*(nos-PhiC31.NLS;)*X*; *P*(carryP)*attP40(II)* and *y*^*1*^*w*\**P*(nos-PhiC31/int.NLS)*X*; *P*(carryP)*attP2(III)* were gifts from the BestGene Inc. *phm22-Gal4*, *spok-Gal4/TM6 Tb* and *spok-Switch-Gal4* were kind gifts from Michael O’Connor’s lab. Stocks were maintained on a cornmeal diet unless otherwise specified.

### Plasmids used for generating transgenic lines

We used the following plasmids that were available from Addgene: pAct:Cas9 (62209) for generating spokHs9, pAct:dCas9-VP64 (78901) for generating spok64 and spokd, pWalium20-10xUAS-3xFlag-dCas9-VPR (78897) for generating spokVPR, pP(ELAV-GeneSwitch) (83957) for generating spokGSa, pBPGUw (17575) for generating gG.Cas9 collection, pUC19 (50005), pCFD3 (49410), pCFD4 (49411) and pCFD5 (73914) (Ornitz et al., 1991; Wang et al., 2012; Port et al., 2014; Lin et al., 2015; Chavez et al., 2016; Port and Bullock, 2016). pAct:FokI-dCas9 (62211) for generatings spokd was a kind gift from Simon Bullock’s lab. pVasa.Cas9 (1340) was obtained from Drosophila Genomics Resources Center for generating spokDm9. All PCR reactions were performed using NEB Q5 high fidelity DNA Polymerase (NEB M0491S) and purified by HighPrep™ PCR reagent from MagBio (AC-60005) following manufacturer’s protocol. All cloning steps were based on the Gibson reaction (Gibson et al., 2009).

### Generating the general Gateway Cas9 (gG-Cas9) collection

The gG-Cas9 collection is based on the pBPGUw plasmid, which we modified to produce different Cas9 versions. This vector contains a Gateway Cassette, a synthetic core promoter and a Gal4-coding sequence (Pfeiffer et al., 2008; Pfeiffer et al., 2010). The pBPGUw backbone was amplified to remove the Gal4 sequence and combined with the different Cas9 versions amplified from corresponding Addgene or DGRC plasmids mentioned above using Gibson reaction (Figure 9, Table S3). Constructs were then transformed into competent DH5α cells and validated by Sanger sequencing.

### Generating the prothoracic gland-specific Cas9 collection (PG-Cas9)

To generate different PG-Cas9 constructs, we used PhiC31 vectors from the above-described gG-Cas9 collection. Vector backbones were amplified via PCR and fused with a 1.45kb fragment containing the *spok* regulatory region amplified from pCRII-TOPO Spok plasmid (a kind gift from Michael O’Connor) via the Gibson reaction (Table S3). Constructs were then transformed into competent DH5α cells and validated by Sanger sequencing.

### Generating prothoracic gland-specific gRNA plasmids (PG-gRNA)

PG-specific gRNA plasmids were generated based on the pCFD5 plasmids that utilize tRNA-flanked gRNAs (Port and Bullock, 2016). We amplified the pCFD5 backbone via PCR and fused the 1.45kb *spok* regulatory region obtained from the CRII-TOPO Spok plasmid (a kind gift from Michael O’Connor). To ensure proper processing of the Pol II-derived transcript, we added either an HV or HDV ribozyme-containing region to the 3’ end, which was then amplified together with tRNA-gRNA duplexes (Figure 9A, Table S3). These fragments were cloned together via Gibson reactions, transformed into DH5α, and validated by Sanger sequencing.

### gRNA selection and cloning

Target gene sequences were obtained from FlyBase and analyzed for optimal target gRNA sites by selecting sequences that showed consensus between two programs, namely “CRISPR optimal target finder” (http://tools.flycrispr.molbio.wisc.edu/targetFinder/) and “Harvard CRISPR gRNA design tool” (http://www.flyrnai.org/crispr/). Optimal target sites were then confirmed by sequencing the loci from genomic DNA extracted from corresponding fly lines we used for plasmid injection. The pCFD5 and PG-gRNA plasmids were pre-digested with BbsI (NEB R3539S) and fused to appropriate gRNA-containing PCR fragments via the Gibson reaction, followed by Sanger sequencing.

### Data availability statement

Fly strains and plasmids made by us are available upon request. The authors affirm that all data necessary for confirming the conclusions of the article are contained within the article, figures, and tables.

### Embryo injections

Plasmids were prepared by Qiagen plasmid midiprep (#12145) and eluted in nuclease-free water (Ambion Life Technologies AM9939) at a concentration of 500-600 ng/μl. Embryo injection was performed either at the University of Alberta or via GenetiVision Corporation following standard procedures (Fish et al., 2007). 300-500 embryos per constructs were injected into *y*^*1*^*w*\**P*(nos-PhiC31.NLS;)*X;P*(carryP)*attP2(II)* or *y*^*1*^*w*\**P* (nos-phiC31/int.NLS)*X;P*(carryP)*attP2(III)* (for CRISPR/Cas9 constructs) or Bloomington #25709, #25710 (for gRNA constructs). Surviving adults were backcrossed to w^1118^ and screened for transformants.

### Survival, 20E-rescue and GeneSwitch studies

Experiments were performed at 25°C and 60-70% humidity. For any fly-based experiments, stocks were reared on NutriFly media starting two generations prior to the actual experiment. Nutrifly (Diamed) is based on the Bloomington Drosophila Stock center formulation (https://bdsc.indiana.edu/information/recipes/bloomfood.html). For egg collections, flies were allowed to lay eggs for 3x 1-hour in order to reduce egg retention and minimize the presence of old embryos. Embryos were then collected in one-hour intervals, counted and transferred to vials containing appropriate media. Larval survival was scored at every stage. At least three independent crosses (~ three biological replicates) were carried out per experimental condition. For 20OH-ecdysone (20E)-supplemented media, the final concentration was 0.33 mg/ml. To activate Cas9-GS (spokGS and spokGSa), mifepristone/RU-486 (Sigma M8046) was used at a final concentration of 100 μg/mL, in line with other studies (Osterwalder et al., 2001; Landis et al., 2015).

### DNA extraction from ring glands

To analyze mutagenesis efficiency in the prothoracic gland (PG) conditional CRISPR, 15-20 ring glands were hand-dissected in collection buffer (10 mM Tris-HCl pH 8.2, 25 mM NaCl, 1 mM EDTA, 0.2% v/v Triton X-100 and 200 μg/mL proteinase K (AM2546)) and incubated for 40 min at 37°C before heat-inactivating proteinase K at 95°C for 5 min. The target region was amplified from the extracted genomic PG DNA via PCR and cloned into the pUC19 vector (NewEngland Biolabs N3041S), which was pre-digested with EcoRI and XbaI. Products were transformed into DH5α competent cells and colonies were randomly selected for Sanger sequencing.

### Immunostains

3^rd^ instar larvae were collected 40-42 hour after the L2/L3 molt and brain-ring gland complexes (BRGC) were dissected and collected in 1xPBS. Samples were fixed in 1x PBS 4% formaldehyde (ThermoFisher 28906) for 20 min at room temperature (RT) before being washed in 1x PBS 0.3% Triton (Sigma T9284) (PBS3T) for 3x 10 min. Samples were blocked at RT for one hour in blocking solution (1x PBS3T 5% normal goat serum (Abcam ab138478)) and then incubated in primary antibody dilution buffer (antibody diluted in 1x PBS3T and 1% BSA) overnight at 4°C with gentle shaking. Samples were then washed in 1x PBS3T for three times with 10 min each, incubated in secondary antibody dilution buffer for 1 hour at room temperature, washed in 1x PBS3T and 1:50,000 DAPI (Cell Signaling 4083) for three times. Samples were mounted in Vectashield mounting medium (VECTH1000). Pictures were then taken using Nikon eclipse 80i confocal C2+ camera. To stain for Cas9, we used anti-CRISPR-Cas9 mouse monoclonal primary antibody (Abcam ab191468) at a ratio of 1:1000 and Goat anti-Mouse IgG H&L Alexa Fluor 555 secondary antibody (Abcam ab150114) was used at a ratio 1:2000.

### *Ex vivo*-culturing and transfection of ring glands

BRGC were dissected from transgenic L3 larvae just after the L2/L3 molt, transferred to culture medium (Schneider insect medium with 10% heat inactivated FBS, 1% streptomycin-penicillin, 10 μg/ml insulin and 2 μg/ml ecdysone), and incubated at 25°C. Larvae carried transgenes expressing gRNAs targeting −110 and −482 bp upstream of *dib* as well as −174 and −423 bp upstream of *phm* (relative to TSS), respectively. These conditions efficiently mimic *in vivo* conditions and allow physiological functions to be studied for up to 48 hours (Prithviraj et al., 2012). BRGC were transfected with *spokCas9* vectors using standard Calcium Phosphate-based transfection method for S2 cell culture (Invitrogen K2780-01). 24 hours post transfection, 50 individual ring glands were collected per replicate, dissolved in Trizol, and stored at −80°C for later qPCR analysis.

### RNA extraction, cDNA synthesis and real-time PCR (qPCR)

Larvae were raised on NutriFly media and staged at 42 hours after the L2/L3 molt. 50 individual ring glands were collected per replicate, dissolved in Trizol, and stored at −80°C for later qPCR analysis. RNA was extracted using Qiagen RNeasy extraction kit (74106) and concentrations were measured via the RNA HS assay kit (Invitrogen Q32852) in a Qubit 2.0 (Invitrogen Q10210). Extracted samples were reverse-transcribed via the ABI High capacity cDNA synthesis kit (ThermoFisher 4368814). Synthesized cDNAs were used for qPCR (QuantStudio 6 Flex, Applied Biosystems) using the Luna^®^ Universal qPCR master mix (NEB M3003S). Samples were normalized to rp49 by calculating fold changes via the ΔΔCT method.

---

**Figure S1.**
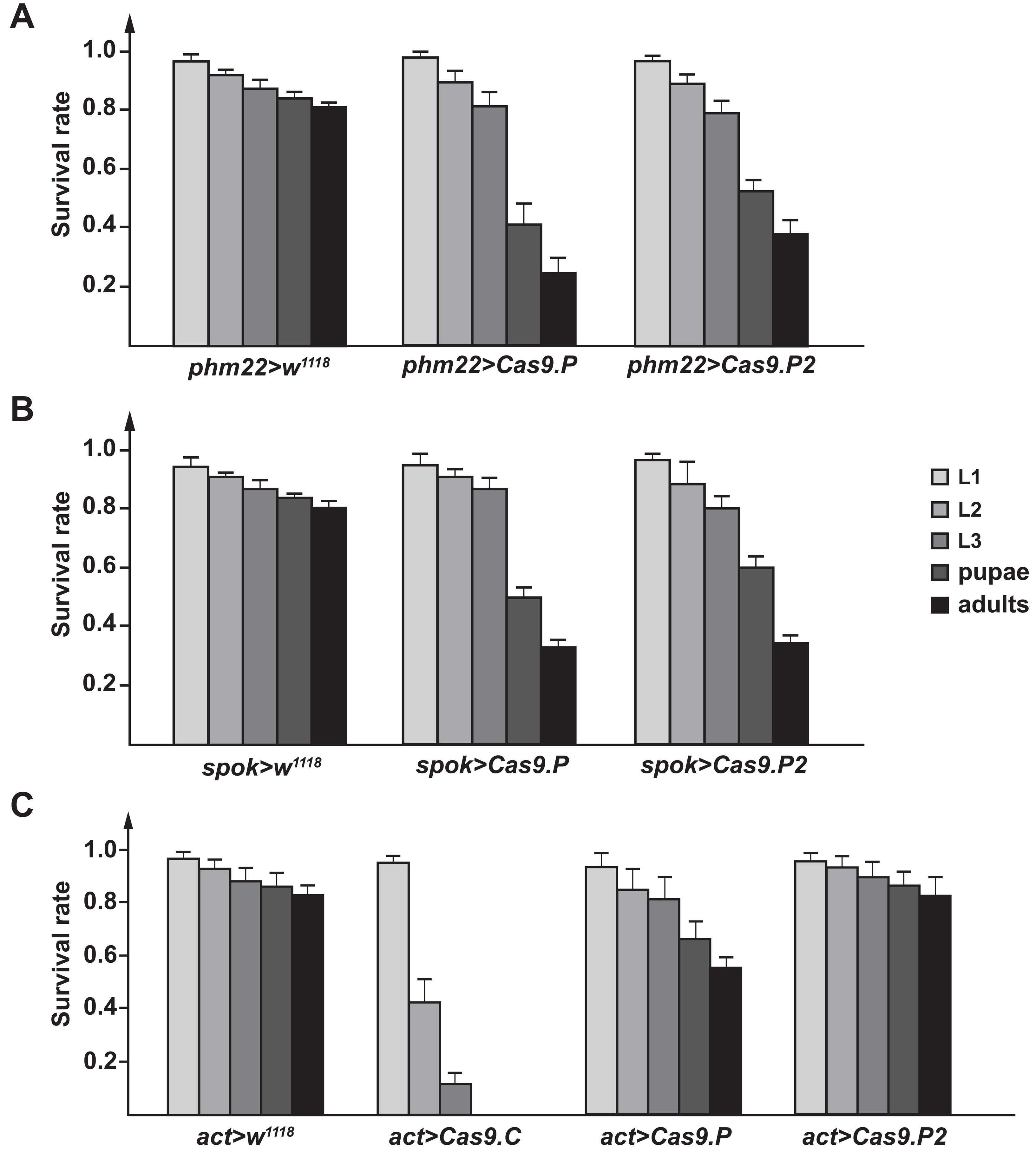
PG-specific and ubiquitous Gal4-driven expression of Cas9 causes lethality. **A.** The survival rates of*phm22>Cas9.P* and*phm22>Cas9.P2* animals. Cas9.P is codon-optimized for Drosophila and Cas9.P2 is codon-optimized for human cells. **B.** The survival rates of *spok>Cas9.P* and *spok>Cas9.P2* animals. **C.** The survival rates of *act>Cas9.C, act> Cas9.P* and *act>Cas9.P2* animals. **A-C.** Data was normalized to the starting number of embryos and error bars represent standard error.

**Figure S2.**
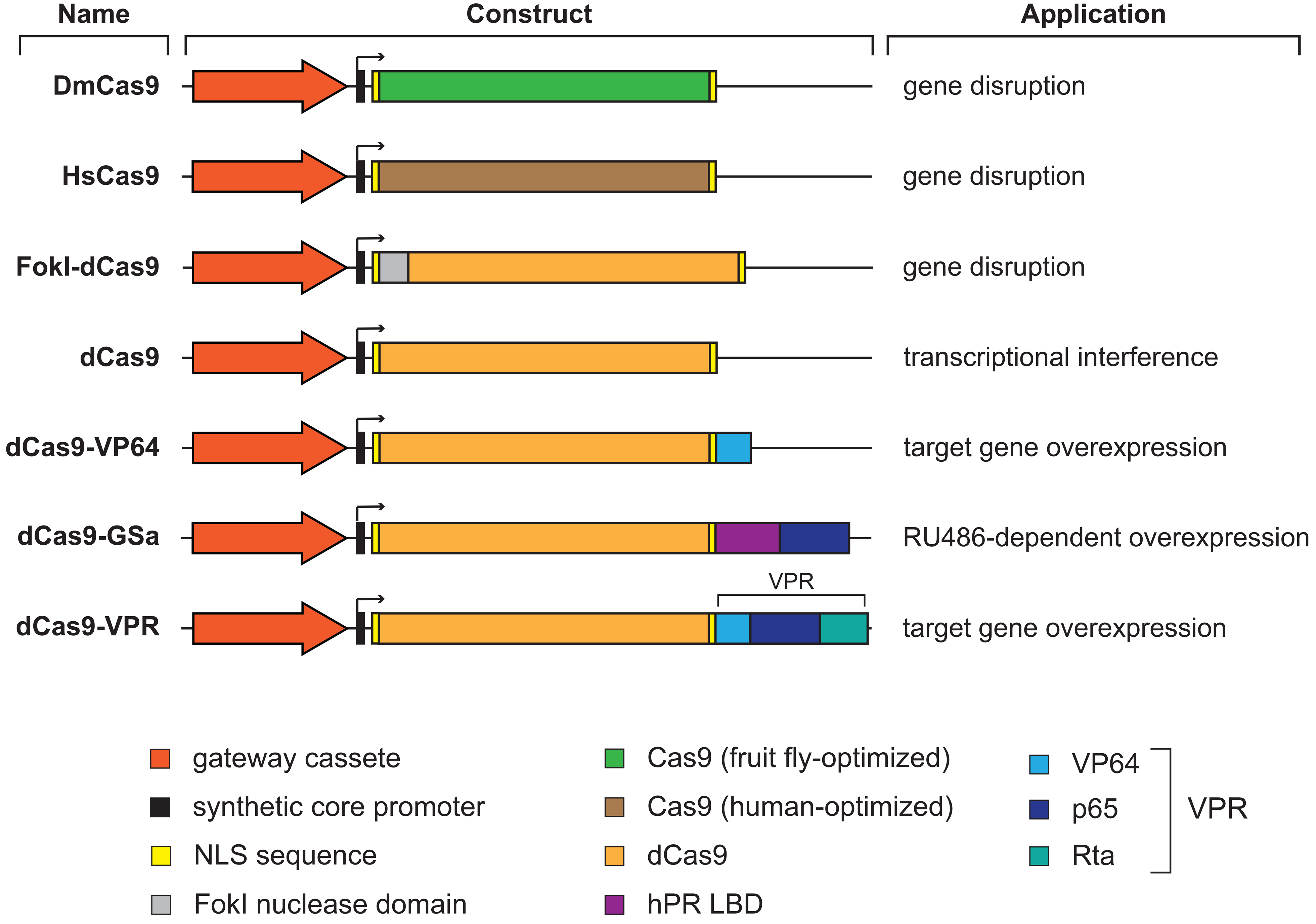
The general Gateway Cas9 (gG-Cas9) vector collection. Each gG-Cas9 vector backbone (not shown) is composed of a mini-*white* gene as a marker, a *PhiC31* integrase-compatible *attB* site, and the *bla* coding sequence to mediate ampicillin resistance. Shown here are the gateway cassette, the Cas9 variant, the regions encoding Nuclear Localization Sequences (NLS), activations domains (VP64, p65 and Rta), the human Progesterone Receptor ligand-binding domain (hPR LBD) and the FokI nuclease domain. The Gateway cassette allows to use LR recombination to insert enhancer/promoter regions to drive tissue-specific Cas9 expression. DmCas9, HsCas9 and FokI.dCas9 can be used to generate somatic mutations. DmCas9 uses a fruit fly codon-optimized Cas9 version, while HsCas9 is optimized for human cells. FokI-dCas9 cuts DNA upon FokI-mediated dimerization followed by FokI cleavage, since dCas (= dead Cas9) is unable to cut DNA (Wah et al., 1997; Guilinger et al., 2014). However, the dCas9 vector can be used to interfere with transcription by guiding Cas9 into the vicinity of transcriptional start sites where it may block the assembly of the pre-initiation complex. dCas9-VP64, dCas9 GSa and dCas9.VPR were designed achieve upregulation of target genes. dCas9-GSa (GeneSwitch activation) encodes a protein where Cas9 is fused to the hPR LBD and p65 domain, allowing activation via RU486.

**Figure S3.**
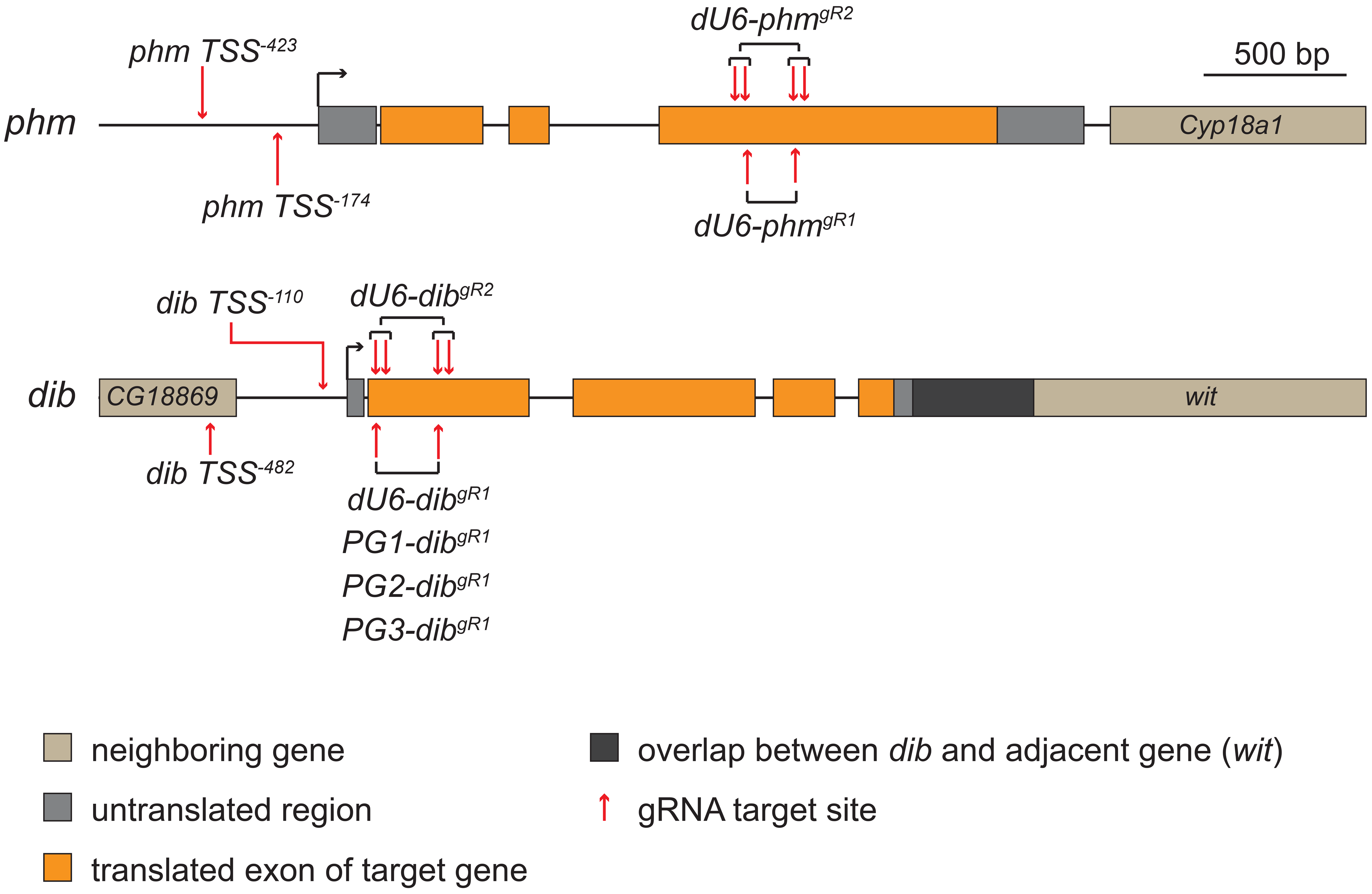
Target sites of gRNAs. We tested two genes that encode enzymes acting as ecdysteroid-synthesizing enzymes in the Drosophila prothoracic gland (PG), *phantom* (*phm*) and *disembodied* (*dib*). gRNAs targeting coding sequence (CDS, orange) were used for somatic disruption via CRISPR. gRNAs that target the upstream region of the of transcription start site (TSS, black arrow) were used either to activate (CRISPRa) or to interfere with (CRISPRi) target gene transcription.

**Table S1. Transgenic gRNA constructs and properties.** In gRNA column, number represents number of gRNAs used to direct Cas9, brackets indicate whether target region is upstream of TSS (= Transcription Start Site) or in the coding region (CDS = Coding Sequence).

**Table S2. Survival data for all CRISPR/CAS9-gRNA crosses.**

**Table S3. Primers used for transgenic lines and vectors.**

